# A minimal functional ZDHHC5 defines distinct requirements for NOD2 binding and *S-*acylation

**DOI:** 10.64898/2026.09.30.755683

**Authors:** Noah R. Martin, Ronan C. Jensen, Gregory D. Fairn

## Abstract

Protein *S-*acylation by ZDHHC5 promotes membrane recruitment of the innate immune receptors NOD1 and NOD2. ZDHHC5 contains an extended cytosolic tail implicated in substrate recruitment, accessory-protein interactions, and enzyme trafficking, but its contribution to NOD1 and NOD2 modification remains unclear. Here, we used mutagenesis and complementation of ZDHHC5-knockout cells to define the regions required for NOD association, *S-*acylation, and plasma membrane localization. Substitution of three *S-*acylated cysteines in the tail preserved ZDHHC5 localization and its ability to restore NOD2 membrane recruitment. Progressive truncation revealed that most of the cytosolic tail was dispensable, with a construct comprising residues 1–230 retaining activity towards both NOD1 and NOD2. Further truncation to amino acids 1-218 impaired NOD *S-*acylation and membrane recruitment without eliminating NOD2 association. Thus, substrate association and productive modification have distinct requirements. C-terminal tail removal also differentially affected accessory-protein association: full-length ZDHHC5 co-immunoprecipitated with both GOLGA7A and GOLGA7B, whereas only GOLGA7B was detectably associated with the 1-230 construct. Together, these findings define a minimal functional ZDHHC5 that supports NOD *S-*acylation and distinguish the requirements for substrate association, catalytic activity, and accessory-protein interactions. They establish that the extended tail is dispensable for NOD modification under the conditions examined, while identifying differential requirements for GOLGA7A and GOLGA7B association with ZDHHC5.

## Introduction

Nucleotide-binding oligomerization domain-containing protein 2 (NOD2) is an intracellular pattern-recognition receptor that detects phosphorylated muramyl dipeptide derived from bacterial peptidoglycan to initiate innate immune signalling^1, 2^. Following activation, NOD2 recruits the kinase RIPK2 to promote downstream NF-κB and MAPK signalling and the production of inflammatory mediators^3^. Beyond this canonical pathway, NOD2 recruits ATG16L1 to sites of bacterial entry to promote autophagy^4^, participates in ER stress-induced inflammatory signalling^5^, and supports antiviral responses through association with the mitochondrial adaptor MAVS^6^. NOD2 function is particularly important at mucosal surfaces, and loss-of-function variants in NOD2 are among the strongest genetic risk factors for Crohn’s disease^7–9^. Although NOD2 is predominantly cytosolic, recruitment to cellular membranes is important for productive signalling, indicating that spatial control of NOD2 is integral to receptor activation^10^.

Protein *S-*acylation is a reversible lipid post-translational modification in which fatty acids, such as palmitate, are attached to cysteine residues through a thioester bond^11, 12^. *S-* acylation regulates diverse aspects of protein function, including membrane association, trafficking, stability, protein-protein interactions, and activity^13, 14^. In mammals, a family of ZDHHC protein acyltransferases catalyzes most S-acylation^14^. Unlike irreversible lipid modifications such as prenylation, the thioester linkage is readily reversible, allowing dynamic regulation of protein localization and function^15, 16^.

In 2019, NOD1 and NOD2 were shown to be substrates of ZDHHC5, and their S-acylation is required for efficient membrane recruitment and antibacterial signalling^17, 18^. Crohn’s disease-associated NOD2 variants with impaired *S-*acylation also exhibit defective membrane recruitment, establishing a functional relationship between this modification and NOD2 activity^17^. More recently, we identified members of the ABHD17 family as NOD1 and NOD2 de-*S-*acylases and showed that inhibition of ABHD17 activity can restore *S-*acylation and membrane localization of several Crohn’s disease-associated NOD2 variants^19^. Together, these findings establish NOD1 and NOD2 *S-*acylation as a reversible regulatory cycle controlled by opposing ZDHHC5 and ABHD17 activities. However, despite the importance of ZDHHC5-mediated *S-*acylation for NOD1 and NOD2 protein function, how ZDHHC5 recognizes these substrates and positions them for lipidation remains poorly understood.

A distinguishing feature of ZDHHC5 is its ∼500-amino-acid cytosolic C-terminal tail. This region is reported to regulate ZDHHC5 localization and interactions with several proteins and contains sites for S-acylation and phosphorylation, as well as trafficking and protein-interaction motifs^20–23^. S-acylation of membrane-proximal cysteines within the tail regulates interactions with the Na+/K+-ATPase and ZDHHC5 accessory proteins^22, 24^, while phosphorylation at more distal sites can alter ZDHHC5 trafficking and substrate access^23^. Recent studies have also implicated AKT- and AMPK-dependent phosphorylation of the ZDHHC5 tail in regulating NOD1 *S-*acylation and signalling^25, 26^. However, whether the extended C-terminal domain is required for ZDHHC5 association with NOD proteins or their subsequent *S-*acylation is unknown. Here, we define the regions of ZDHHC5 required for NOD2 association and modification and identify a minimal ZDHHC5 construct that retains substrate binding, *S-* acyltransferase activity, and selective association with GOLGA7B.

## Results

### ZDHHC5 catalytic activity is required for NOD2 plasma membrane localization

To establish a cellular assay to examine the requirements for ZDHHC5-dependent NOD2 membrane localization, we compared the subcellular distribution of GFP-NOD2 in wild-type and ZDHHC5-knockout HEK293T cells. GFP-NOD2 exhibited a readily detectable plasma membrane pool in wild-type cells, whereas this enrichment was largely absent in ZDHHC5-knockout cells (Fig. 1A, B), consistent with previous studies^17^. Re-expression of wild-type mCherry-ZDHHC5 restored GFP-NOD2 plasma membrane localization, whereas the catalytically inactive C134S mutant did not (Fig. 1C, D). These results show that ZDHHC5 catalytic activity is required for NOD2 plasma membrane enrichment in this system and establish a complementation assay to evaluate ZDHHC5 mutants.

**Figure 1.**
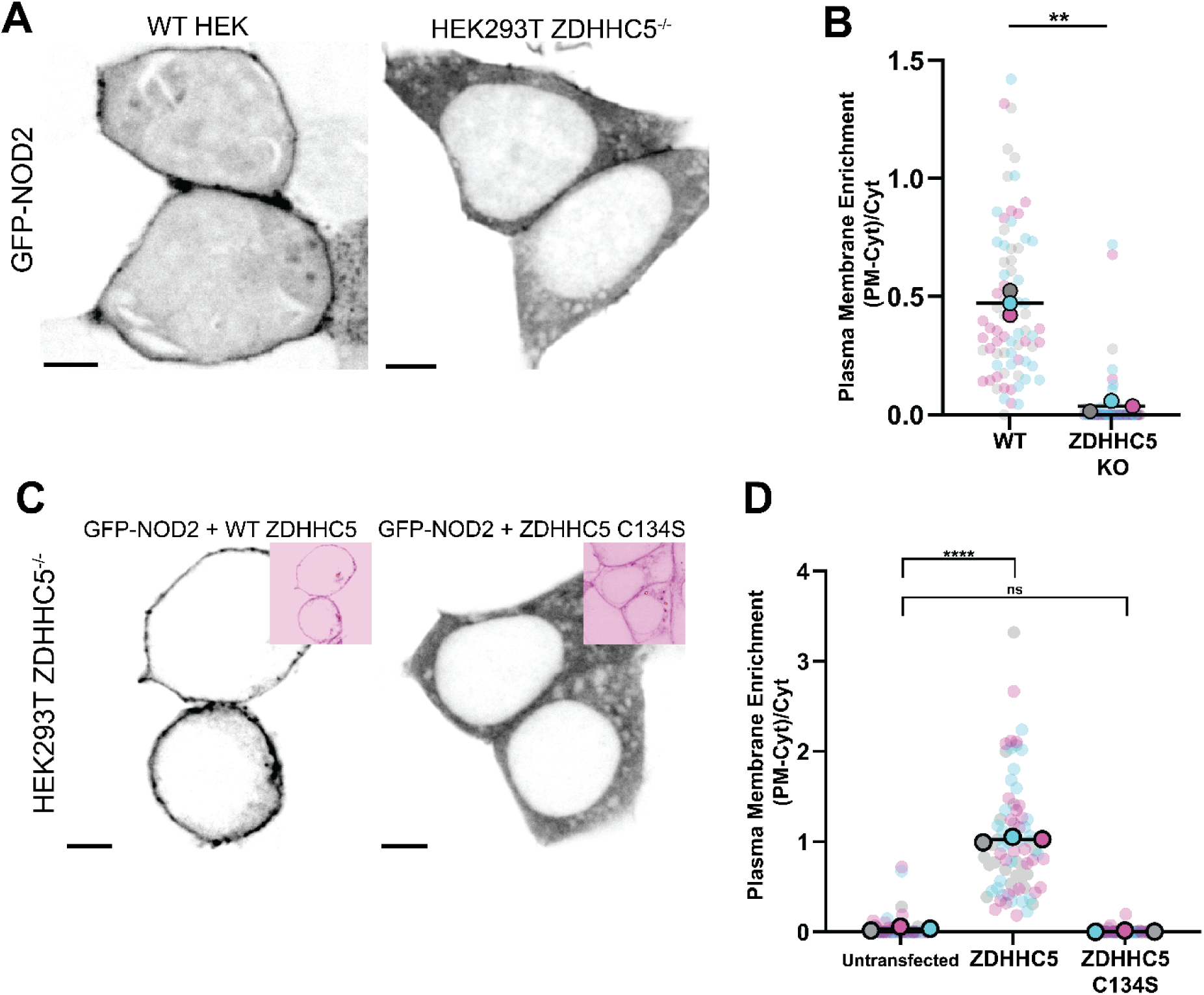
ZDHHC5 catalytic activity is required for NOD2 plasma membrane localization. A) Representative confocal images of wild-type (WT) and ZDHHC5-KO HEK293T cells transiently expressing GFP-NOD2, displayed in inverted grayscale. B) Quantification of GFP-NOD2 plasma membrane localization in WT and ZDHHC5-KO cells, as shown in A. The following data presentation is used throughout: Small dots represent individual cells, and large dots represent the mean of each biological replicate. Colors distinguish the three independent experiments, and black horizontal lines indicate overall means. P values were determined using one-tailed t-tests. *P ≤ 0.05; **P ≤ 0.01; ****P ≤ 0.0001; ns, not significant. C) Representative confocal images of ZDHHC5-KO HEK293T cells transiently co-expressing GFP-NOD2 and either mCherry-ZDHHC5 or the catalytically inactive mCherry-ZDHHC5^C134S^ mutant, as indicated. GFP-NOD2 is displayed in inverted grayscale; mCherry-tagged constructs are shown in the insets in inverted red. D) Quantification of GFP-NOD2 plasma membrane localization under the conditions shown in C. Scale bars, 5 µm. Data are from three independent experiments; colors distinguish independent experiments, and black horizontal lines indicate means. P values were determined by one-way ANOVA followed by Dunnett’s multiple-comparisons test. *P ≤ 0.05; **P ≤ 0.01; ****P ≤ 0.0001; ns, not significant.

### The three S-acylation sites in the ZDHHC5 cytosolic tail are dispensable for NOD2 plasma membrane recruitment

We first examined whether the three S-acylation sites in the membrane-proximal cytosolic tail of ZDHHC5 contribute to its localization or ability to recruit NOD2 to the plasma membrane. We generated a single C245S mutant, a double C236S, C237S mutant, and a triple C236S, C237S, C245S mutant (Fig. 2A). Each construct was transiently expressed in ZDHHC5-knockout cells together with a GFP-tagged version of the pleckstrin homology domain of phospholipase Cδ (PH-PLC), a marker of the plasma membrane due to its ability to bind phosphatidylinositol 4,5-bisphosphate^27^. All three mutants retained prominent plasma membrane localization, as validated through Pearson’s colocalization coefficient^28^ with GFP-PH-PLC, comparable to wild-type mCherry-ZDHHC5 (Fig. 2B, C).

**Figure 2.**
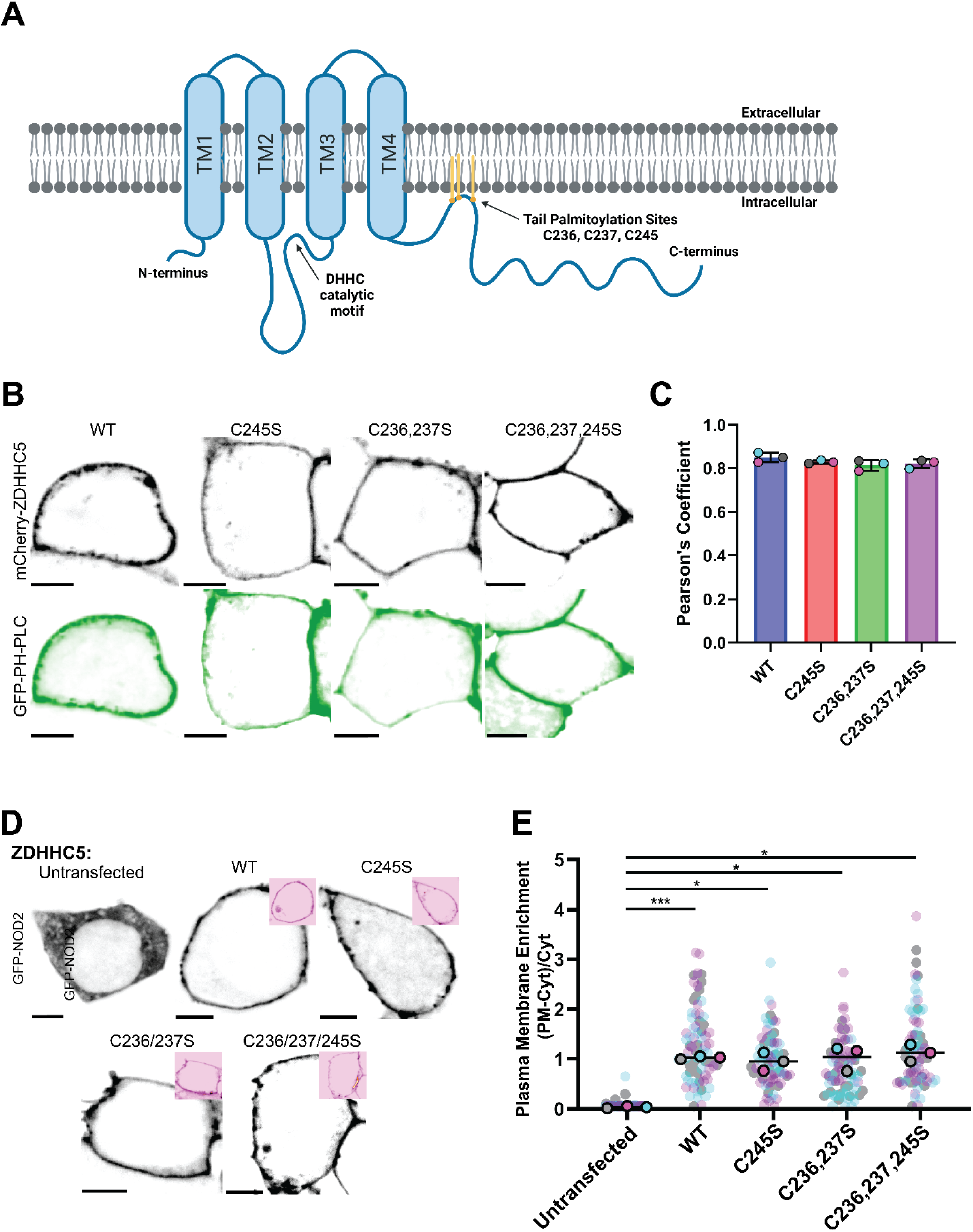
*S-*acylation sites in the ZDHHC5 cytosolic tail are dispensable for NOD2 plasma membrane localization. A) Schematic of ZDHHC5 showing the three cytosolic tail *S-*acylation sites (Cys236, Cys237, and Cys245) targeted for cysteine-to-serine substitution. B) Representative confocal images of ZDHHC5-KO HEK293T cells transiently co-expressing the indicated mCherry-ZDHHC5 constructs (inverted grayscale) and GFP–PH-PLCδ1 (inverted green) as a plasma membrane marker. C) Colocalization of mCherry-ZDHHC5 constructs with GFP–PH-PLCδ1, quantified using Pearson’s correlation coefficient in ImageJ with the BIOP JACoP plugin. Only significant differences are indicated. D) Representative confocal images of ZDHHC5-KO HEK293T cells transiently co-expressing GFP-NOD2 (inverted grayscale) and the indicated mCherry-ZDHHC5 constructs (inverted red insets). E) Quantification of GFP-NOD2 plasma membrane localization under the conditions shown in D. Scale bars, 5 µm. Small dots represent individual cells, and large dots represent the mean of each biological replicate. Colors distinguish the three independent experiments, and black horizontal lines indicate overall means. P values for C and E were determined by one-way ANOVA followed by Dunnett’s multiple-comparisons test. *P ≤ 0.05; ***P ≤ 0.001.

Prominent plasma membrane localization alone does not necessarily mean these constructs are functional or can modify NOD2. Thus, we next assessed whether these substitutions affected NOD2 localization. Each mutant restored GFP-NOD2 plasma membrane enrichment in ZDHHC5-knockout cells, including the triple mutant lacking all three S-acylation sites (Fig. 2D, E). Thus, these cysteine residues are dispensable for both ZDHHC5 plasma membrane localization and its ability to support NOD2 plasma membrane recruitment under the conditions examined.

### Most of the ZDHHC5 cytosolic tail is dispensable for NOD2 plasma membrane recruitment

A previous study generated a series of deletion mutants to identify the region of the cytosolic tail responsible for binding phospholemman^20^. We used a similar approach to determine if a particular region of the extended ZDHHC5 cytosolic tail is required for NOD2 membrane recruitment. We generated a series of truncation constructs designated 527X, 335X, 275X, and 219X (Fig. 3A)^20^. Again, all four constructs retained plasma membrane localization when expressed in ZDHHC5-knockout cells, as assessed by microscopy and colocalization with GFP-PH-PLC (Fig. 3B, C). Therefore, removal of most of the cytosolic tail did not prevent ZDHHC5 from reaching the plasma membrane.

**Figure 3.**
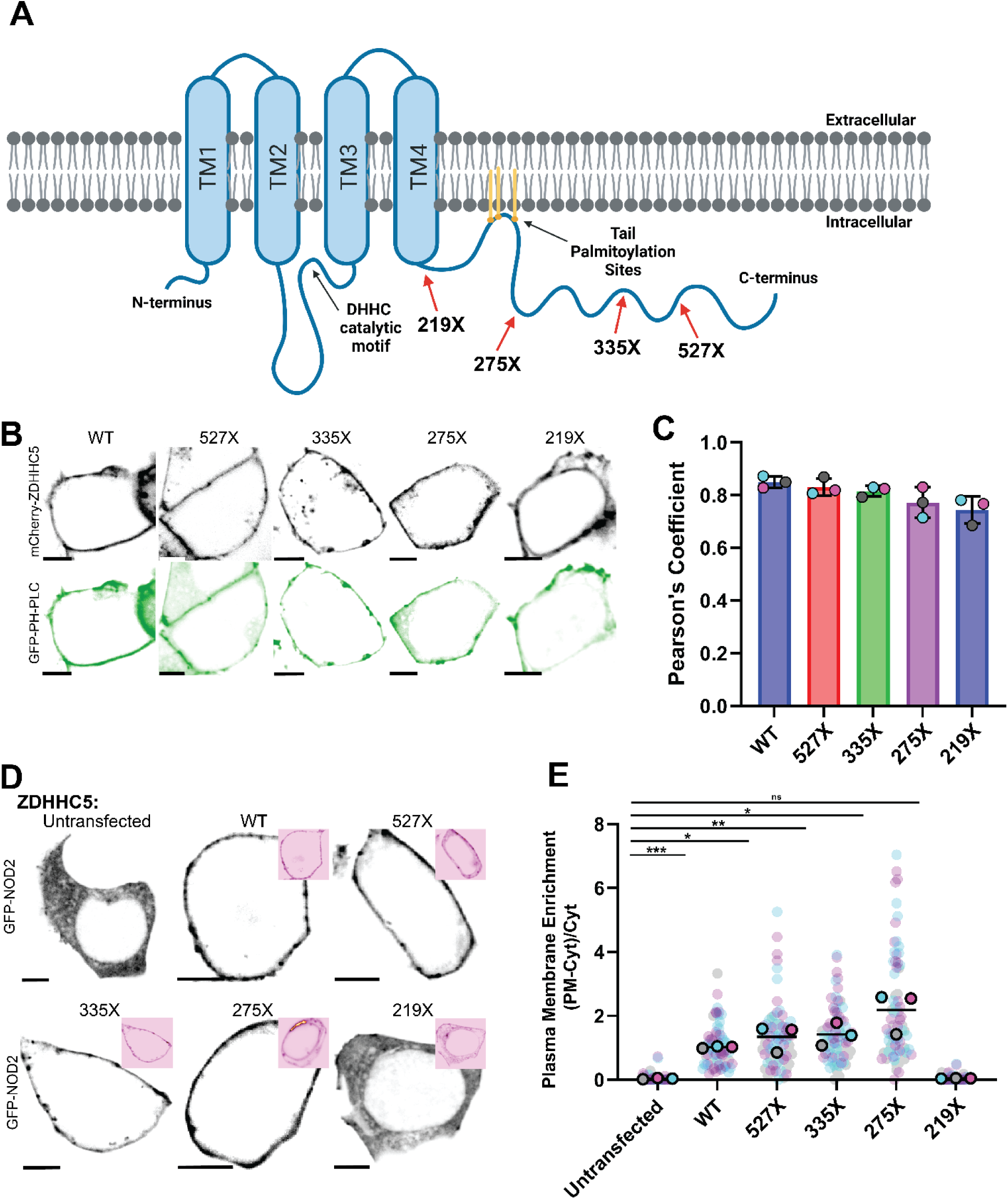
Most of the ZDHHC5 cytosolic tail is dispensable for NOD2 plasma membrane localization. A) Schematic of ZDHHC5 showing the positions of four cytosolic tail truncations. B) Representative confocal images of ZDHHC5-KO HEK293T cells transiently co-expressing the indicated mCherry-ZDHHC5 constructs (inverted grayscale) and GFP–PH-PLCδ1 (inverted green) as a plasma membrane marker. C) Colocalization of mCherry-ZDHHC5 constructs with GFP–PH-PLCδ1, quantified using Pearson’s correlation coefficient in ImageJ with the BIOP JACoP plugin. Only significant differences are indicated. D) Representative confocal images of ZDHHC5-KO HEK293T cells transiently co-expressing GFP-NOD2 (inverted grayscale) and the indicated mCherry-ZDHHC5 constructs (inverted red insets). E) Quantification of GFP-NOD2 plasma membrane localization under the conditions shown in D. Scale bars, 5 µm. Small dots represent individual cells, and large dots represent the mean of each biological replicate. Colors distinguish the three independent experiments, and black horizontal lines indicate overall means. P values for C and E were determined by one-way ANOVA followed by Dunnett’s multiple-comparisons test. *P ≤ 0.05; **P ≤ 0.01; ***P ≤ 0.001; ns, not significant.

The 527X, 335X, and 275X truncation constructs each restored GFP-NOD2 plasma membrane enrichment (Fig. 3D, E). In contrast, the 219X construct failed to restore NOD2 membrane localization despite retaining its own plasma membrane localization. These findings indicate that most of the ZDHHC5 cytosolic tail is dispensable for NOD2 recruitment and identify residues 218-274 as containing determinants required for this activity.

### ZDHHC5 amino acids 1-230 is sufficient to support NOD2 plasma membrane localization

Since the results in Figure 3 demonstrated that the region between 218 and 274 was critical for the ability of ZDHHC5 to localize NOD2 to the plasma membrane, we refined the functional boundary using additional truncations designated 261X, 251X, 241X, and 231X (Fig. 4A). The 251X, 241X, and 231X constructs retained substantial colocalization with GFP-PH-PLC whereas the 261X construct exhibited a reduced localization to the PM; Pearson’s correlation coefficient of ∼0.4 compared to ∼0.8 for wild-type and the other mutants (Fig. 4B, C). Yet despite this difference in distribution, all four constructs supported GFP-NOD2 plasma membrane enrichment in ZDHHC5-knockout cells (Fig. 4D, E). The shortest active construct, ZDHHC5-231X, restored NOD2 membrane localization, whereas ZDHHC5-219X remained ineffective. Thus, amino acids 1-230 define the minimal functional construct within the truncation series tested. Its activity also demonstrates that the downstream tail region containing Cys236, Cys237, and Cys245 is not required for NOD2 plasma membrane recruitment, consistent with the cysteine-substitution experiments.

**Figure 4.**
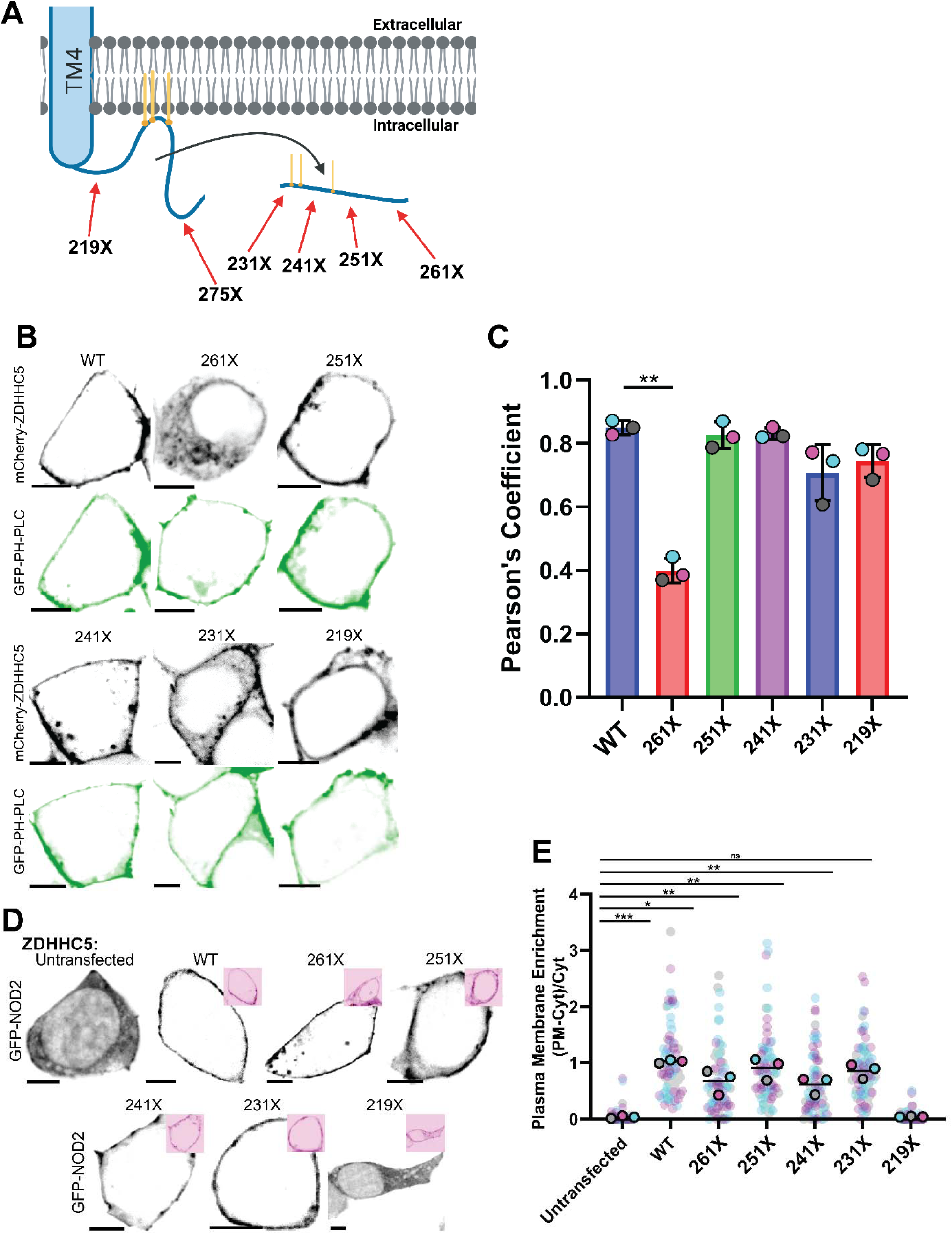
ZDHHC5 1-230 is sufficient to support NOD2 plasma membrane localization. A) Enlarged schematic of the membrane-proximal ZDHHC5 cytosolic tail showing the positions of four additional truncations. B) Representative confocal images of ZDHHC5-knockout (KO) HEK293T cells transiently co-expressing the indicated mCherry-ZDHHC5 constructs (inverted grayscale) and GFP–PH-PLCδ1 (inverted green) as a plasma membrane marker. C) Colocalization of mCherry-ZDHHC5 constructs with GFP–PH-PLCδ1, quantified using Pearson’s correlation coefficient in ImageJ with the BIOP JACoP plugin. Only significant differences are indicated. D) Representative confocal images of ZDHHC5-KO HEK293T cells transiently co-expressing GFP-NOD2 (inverted grayscale) and the indicated mCherry-ZDHHC5 constructs (inverted red insets). E) Quantification of GFP-NOD2 plasma membrane localization under the conditions shown in D. Scale bars, 5 µm. Small dots represent individual cells, and large dots represent the mean of each biological replicate. Colors distinguish the three independent experiments, and black horizontal lines indicate overall means. P values for C and E were determined by one-way ANOVA followed by Dunnett’s multiple-comparisons test. *P ≤ 0.05; **P ≤ 0.01; ***P ≤ 0.001; ns, not significant.

### ZDHHC5 truncation mutants separate the ability to interact with NOD2 from S-acylation

The failure of 219X to restore NOD2 plasma membrane localization could reflect impaired substrate association or an inability to modify an associated substrate. To distinguish these possibilities, we co-expressed FLAG-NOD2 with the mCherry-tagged ZDHHC5 constructs and examined their association by co-immunoprecipitation. FLAG-NOD2 co-immunoprecipitated with all truncation constructs tested, including 219X, and with the catalytically inactive C134S mutant (Fig. 5A). Consequently, the loss of NOD2 membrane recruitment by 219X and the ZDHHC5^C134S^ mutant is not due to loss of detectable NOD2 binding.

**Figure 5.**
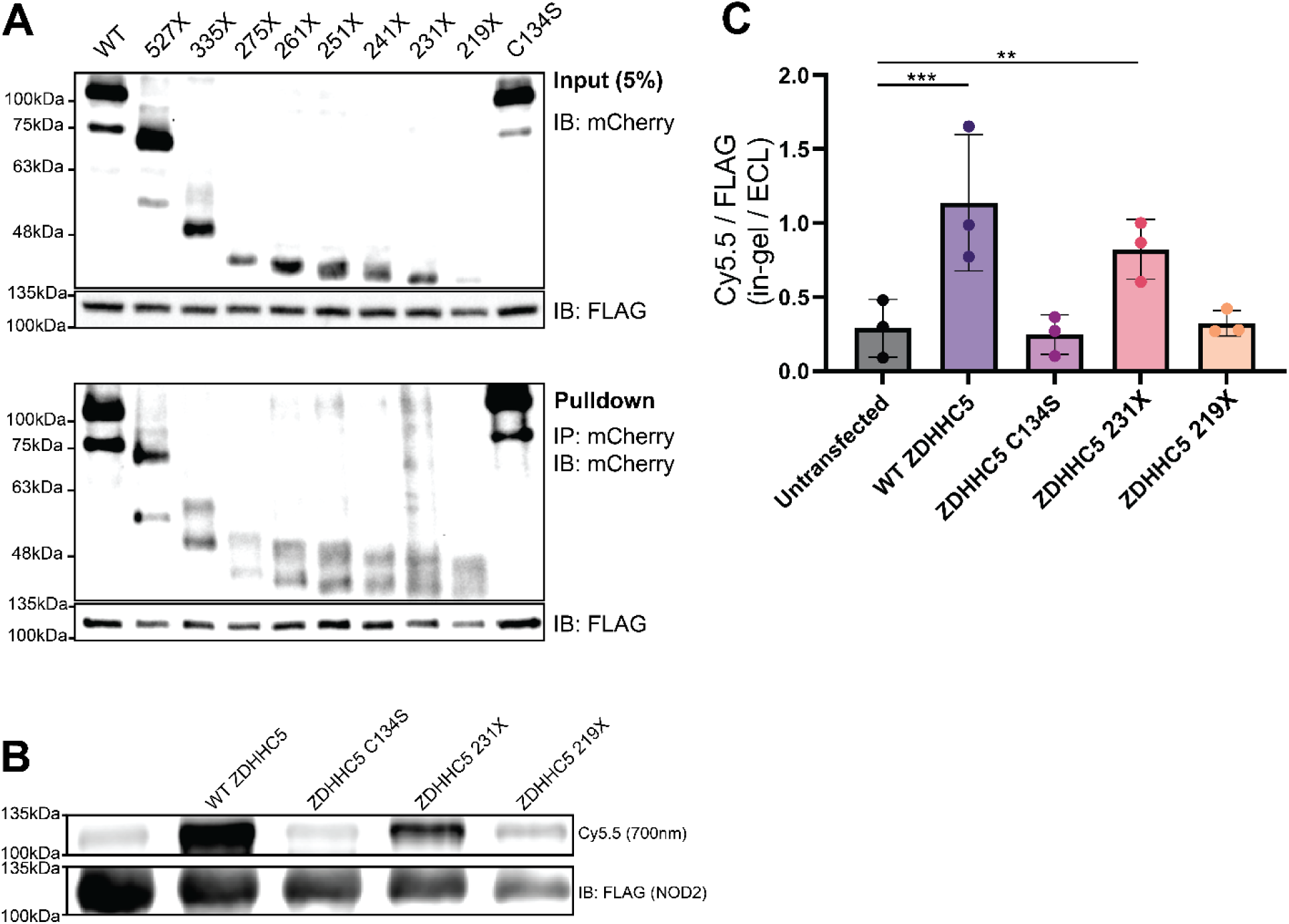
ZDHHC5 truncation mutants distinguish NOD2 association from *S-*acylation. A) Co-immunoprecipitation of FLAG-NOD2 with mCherry-ZDHHC5 constructs transiently co-expressed in HEK293T cells. mCherry-tagged proteins were isolated using RFP-Catcher agarose beads, and associated FLAG-NOD2 was detected by immunoblotting. B) Detection of NOD2 *S-* acylation by metabolic labeling and click chemistry. ZDHHC5-KO HEK293T cells transiently co-expressing FLAG-NOD2 and the indicated mCherry-ZDHHC5 constructs were incubated with the alkyne-functionalized fatty acid analog 15-HDYA. FLAG-NOD2 was immunoprecipitated using anti-FLAG agarose beads and subjected to on-bead click chemistry. Incorporation of 15-HDYA was detected by Cy5.5 in-gel fluorescence using the 700-nm channel. Following membrane transfer, FLAG-NOD2 was detected by immunoblotting with chemiluminescent detection. C) Quantification of FLAG-NOD2 S-acylation under the conditions shown in B, calculated as the ratio of Cy5.5 fluorescence to the corresponding FLAG-NOD2 immunoblot signal. Blots are representative of three independent experiments; quantification includes all three experiments. P values for C were determined by one-way ANOVA followed by Dunnett’s multiple-comparisons test. **P ≤ 0.01; ***P ≤ 0.001; ns, not significant.

We then directly assessed NOD2 acylation by metabolic labelling with the clickable fatty acid analogue 15-hexadecynoic acid (15-HDYA). After recovering FLAG-NOD2 with anti-FLAG beads, we detected the attached 15-HDYA using copper-catalyzed azide-alkyne cycloaddition (CuAAC) to attach Azide-Cy5.5 and assess in-gel fluorescence. Wild-type ZDHHC5 and ZDHHC5-231X significantly increased NOD2 labelling relative to the control cells lacking ZDHHC5. In contrast, expression of ZDHHC5^C134S^ or ZDHHC5-219X in the ZDHHC5^KO^ cells yielded signals comparable to control levels (Fig. 5B, C). These biochemical findings agree with the localization assays and demonstrate that ZDHHC5-231X retains activity towards NOD2. Together, the association and *S-*acylation experiments define distinct requirements for NOD2 association and modification: 219X retains detectable substrate association but does not support efficient NOD2 *S-*acylation and, as a result, fails to support NOD2 localization to the plasma membrane.

### ZDHHC5 1-230 *S-*acylates NOD1

To determine whether the activity of this minimal ZDHHC5 construct extends to NOD1, we expressed GFP-NOD1 together with wild-type or mutant ZDHHC5 in ZDHHC5^KO^ cells. Both wild-type ZDHHC5 and ZDHHC5-231X promoted GFP-NOD1 plasma membrane localization, whereas C134S and 219X did not produce comparable membrane enrichment (Fig. 6A). Metabolic labelling with 15-HDYA and CuAAC reaction with Azide-Cy5.5 similarly showed that wild-type ZDHHC5 and ZDHHC5-231X significantly increased FLAG-NOD1 acylation relative to the control lacking re-expressed ZDHHC5. In contrast, signals obtained with the C134S and 219X mutants remained near control levels (Fig. 6B, C). Thus, ZDHHC5 amino acids 1-230 support the *S-*acylation and plasma membrane localization of both NOD1 and NOD2.

**Figure 6.**
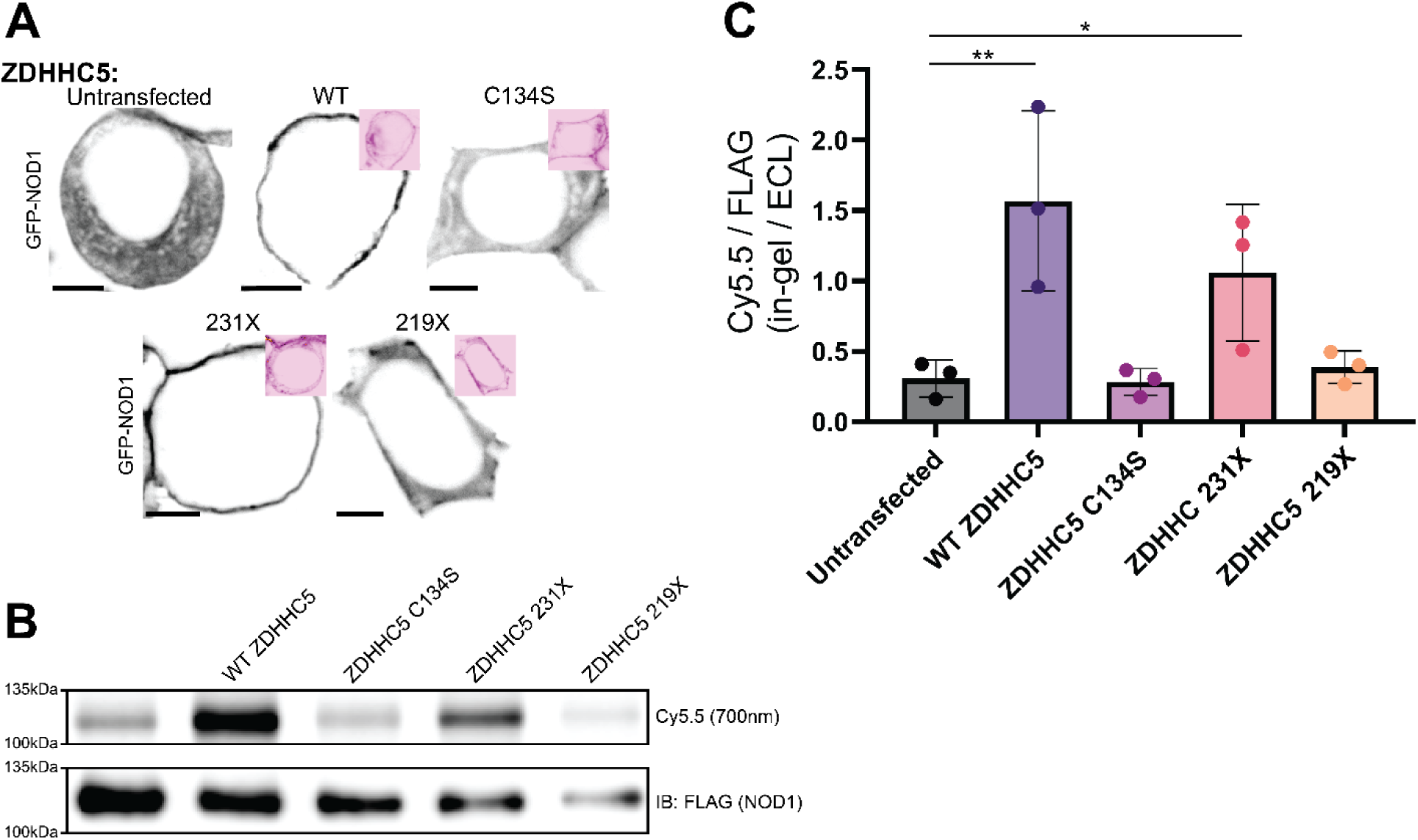
ZDHHC5 1-230 supports NOD1 S-acylation and plasma membrane localization. A) Representative confocal images of ZDHHC5-KO HEK293T cells transiently co-expressing GFP-NOD1 (inverted grayscale) and the indicated mCherry-ZDHHC5 constructs, including the catalytically inactive C134S control (inverted red insets). Scale bars, 5 µm. B) Detection of NOD1 S-acylation by metabolic labeling and click chemistry. ZDHHC5-KO HEK293T cells transiently co-expressing FLAG-NOD1 and the indicated mCherry-ZDHHC5 constructs were incubated with the alkyne-functionalized fatty acid analog 15-HDYA. FLAG-NOD1 was immunoprecipitated using anti-FLAG agarose beads and subjected to on-bead click chemistry. Incorporation of 15-HDYA was detected by Cy5.5 in-gel fluorescence in the 700-nm channel. Following membrane transfer, FLAG-NOD1 was detected by immunoblotting with chemiluminescent detection. C) Quantification of FLAG-NOD1 S-acylation under the conditions shown in B, calculated as the ratio of Cy5.5 fluorescence to the corresponding FLAG-NOD1 immunoblot signal. Images and blots are representative of three independent experiments; quantification includes all three experiments. P values for C were determined by one-way ANOVA followed by Dunnett’s multiple-comparisons test. *P ≤ 0.05; **P ≤ 0.01; ns, not significant.

### ZDHHC5 1-230 retains association with GOLGA7B but not GOLGA7A

ZDHHC5 associates with the related accessory proteins GOLGA7A and GOLGA7B, which can influence its cellular functions. Evidence suggests that *S-*acylated GOLGA7B stabilizes ZDHHC5 at the plasma membrane, providing a mechanism through which an accessory protein can regulate its localization^22, 29^. We therefore asked whether the minimal ZDHHC5 construct retains association with these proteins.

FLAG-GOLGA7A and FLAG-GOLGA7B were each recovered in co-immunoprecipitates of full-length mCherry-ZDHHC5 (Fig. 7A). In contrast, FLAG-GOLGA7B, but not FLAG-GOLGA7A, was recovered with ZDHHC5-231X under the conditions tested (Fig. 7B). Thus, removal of most of the cytosolic tail preserves association with GOLGA7B while differentially affecting association with GOLGA7A.

**Figure 7.**
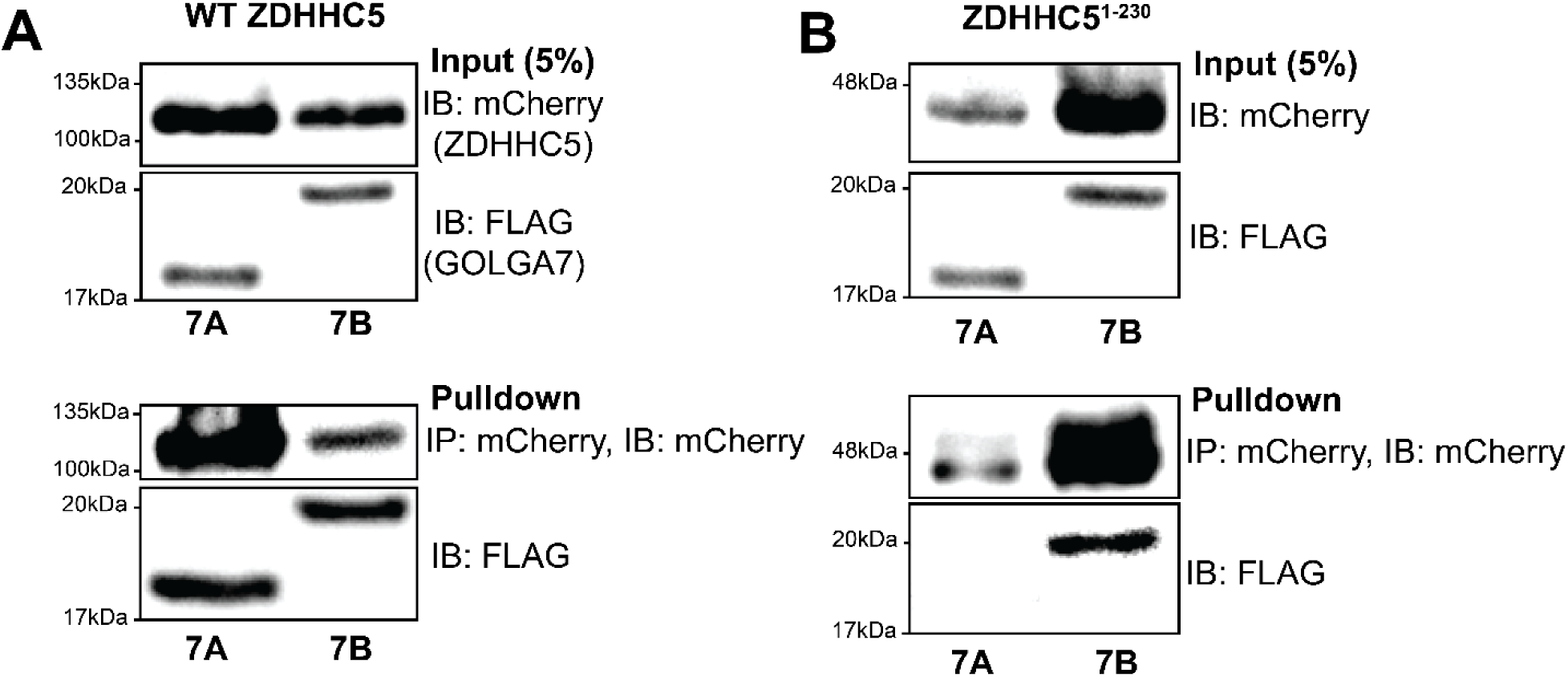
ZDHHC5 1-230 retains association with GOLGA7B but not GOLGA7A. A) Co-immunoprecipitation of FLAG-GOLGA7A or FLAG-GOLGA7B with full-length mCherry-ZDHHC5 transiently co-expressed in HEK293T cells. mCherry-ZDHHC5 was isolated using RFP-Catcher agarose beads, and associated FLAG-tagged proteins were detected by immunoblotting. B) Co-immunoprecipitation of FLAG-GOLGA7A or FLAG-GOLGA7B with mCherry-ZDHHC5 1-230, performed as described in A.

## Discussion

Our findings define a minimal ZDHHC5 protein that supports NOD1 and NOD2 *S-*acylation and plasma membrane localization. Substitution of the three *S-*acylated cysteines in the cytosolic tail of ZDHHC5 did not impair NOD2 membrane recruitment, and progressive truncation showed that most of the tail is dispensable for its ability to modify NOD1 and NOD2. ZDHHC5^1–230^ retained activity toward both NOD proteins, whereas further truncation to 219X abolished efficient modification without eliminating NOD2 association, as measured by co-immunoprecipitation. The ZDHHC5^1–218^ shorter truncation interrupts the conserved TTxE motif (aa 216-219) immediately following the fourth transmembrane domain. Structural and mutational analyses of ZDHHC20 have shown that this motif participates in intramolecular interactions that organize the catalytic region and support enzymatic activity^30^. Disruption of this motif plausibly explains the inactivity of 219X. Together, these findings distinguish the requirements for NOD2 association from those for productive S-acylation and demonstrate that most of the extended cytosolic tail is unnecessary for NOD modification under the conditions examined.

Removal of the cytosolic tail differentially affected ZDHHC5 association with the accessory proteins GOLGA7A and GOLGA7B. Whereas both proteins co-immunoprecipitated with full-length ZDHHC5, only GOLGA7B was detectably recovered with the ZDHHC5^1-230^ construct. Recent structural and mutational analyses identified conserved determinants of ZDHHC5– GOLGA7A complex assembly, including the ZDHHC5 RNYR motif and the GOLGA7A RDYS motif, with the latter positioned against the zinc-binding region of ZDHHC5^31^. Although the resolution limited assignment of some individual side-chain contacts, mutagenesis studies supported these interfaces. The corresponding ZDHHC5 determinants are retained in 1-230 mutants, suggesting that their presence alone is insufficient to maintain stable association to support GOLGA7A detection in the immunoprecipitation experiments. Additional contacts or conformational effects involving the membrane-proximal tail may therefore contribute differently to assembly of the two complexes.

The largely dispensable nature of the ZDHHC5 tail for NOD1 and NOD2 *S-*acylation contrasts with its established role in recognizing or recruiting other substrates. GRIP1b binding and *S-*acylation require the last four amino acids of the C-terminus, EISV, which serve as a PDZ-binding motif, illustrating the role of the protein’s extreme distal end in substrate recognition^21^. The previously mentioned studies of phospholemman identified a requirement for the membrane-proximal tail and subsequently mapped the association of the Na /K -ATPase complex to ZDHHC5 residues 233–257^20, 24^. This interaction also provides a mechanism to position the pump-associated phospholemman for modification without requiring most of the C-terminal tail. Peptide-mapping experiments also identified associations of Sodium-Calcium Exchanger 1 with residues 233-267 of ZDHHC5, extending the evidence for this region near transmembrane 4 as a recruitment site for membrane-protein substrates^32^. Together, these observations support substrate-specific requirements for the ZDHHC5 tail, with some substrates relying on additional docking interactions and others accessing the enzyme through determinants retained within its minimal functional core.

Beyond substrate recruitment, the unusually long ZDHHC5 C-terminal tail provides multiple opportunities to regulate where and when *S-*acylation occurs. Fyn-mediated phosphorylation of Tyr533 inhibits association with endocytic machinery and stabilizes ZDHHC5 at the neuronal surface, whereas neuronal activity promotes redistribution of the enzyme to intracellular compartments where it encounters additional substrates^23^. More recently, AMPK and AKT were reported to phosphorylate distinct, partially overlapping sets of tail residues, respectively reducing or promoting ZDHHC5 membrane association and NOD1 modification^25, 26^. These findings suggest that the extended tail allows extracellular and metabolic signals to regulate substrate access by changing enzyme localization. For a plasma membrane-enriched acyltransferase like ZDHHC5, control of surface retention, endocytosis, and endosomal trafficking could be especially important for coordinating modification with receptor activation and membrane recycling. Our results are consistent with a model in which much of the tail provides context-dependent regulation beyond the minimal requirements for NOD *S-* acylation. Its dispensability in our complementation assays therefore does not imply dispensability during physiological signalling; rather, it separates the capacity to modify NOD proteins from the additional mechanisms that may control that activity in space and time.

### Experimental procedures

#### Reagents and Fine Chemicals

X-tremeGENE HP (Milipore-Sigma, Cat#: 6366244001, Lot#: 89878600), Fatty-acid-free bovine serum albumin (BioBasic, Cat#: AR2448, Lot#: P6210010), alkynyl palmitic acid (15-HDYA) (Vector Laboratories, Cat#: CCT-1165, Lot#: 6197B), cOmplete EDTA-free protease inhibitor (Millipore-Sigma, Cat#: 05056489001, Lot#: 86011100), high-glucose Dulbecco’s Modified Eagle’s Medium (Wisent, Cat#: 319-020-CL, Lot#: 319020062), fetal bovine serum (Wisent, Cat#: 080-150, Lot#: 112768), FLAG affinity beads (Sigma-Aldrich, Cat#: A2220, Lot#: 0000472626), THPTA (Vector Laboratories, Cat#: CCT-1010, Lot#: 5987A), CuSO_4_ ( Sigma-Aldrich, Cat#: 1003398912, Lot#: MKCQ4337), sodium ascorbate (300 mM; Jena Bioscience, Cat#: CLK-I005, Lot#: KLI019-004), and Cy5.5 Picolyl Azide (2 mM; Vector Laboratories, Cat#: CCT-1182, Lot#: 1676X)

#### Cell Culture and Transfection

Wild-type (WT) human embryonic kidney (HEK) 293T were purchased from the American Type Culture Collection (ATCC, Manassas, Virginia). HEK293T ZDHHC5 knockout cells were a gift from Dr. Scott Dixon (Department of Biology, Stanford University)^31^. Cell lines were maintained and propagated in high-glucose Dulbecco’s Modified Eagle’s Medium (DMEM) supplemented with 1.5 g/L sodium bicarbonate, L-glutamine, and 10% fetal bovine serum (FBS) in a humidified incubator at 37°C with 5% CO2. We routinely tested cells for *Mycoplasma* using a PCR-based test. HEK293T WT and ZDHHC5 knockout cells were transfected at 80% confluency using X-tremeGENE HP according to the manufacturers protocol. The following day cells were harvested, treated, or fixed for confocal imaging.

#### Constructs

FLAG-NOD1, FLAG-NOD2, and mCherry-ZDHHC5 were inherited from a previous study in the lab^17^. GFP-NOD1 and GFP-NOD2 were gifts from Dr. Charneal Dixon (Department of Pathology, Dalhousie University)^19^.

#### Site-directed Mutagenesis and Truncation Mutants

The large and small scale ZDHHC5 truncations were generated by restriction cloning, primers were used to amplify the region of ZDHHC5 indicated with a stop codon included in the reverse primer, truncated fragments of ZDHHC5 were inserted into the mCherry-C1 vector (TaKaRa Clontech, Cat#: 632524) via restriction cloning. ZDHHC5 Cys-to-Ser mutants (C134S, C245S, C236,237S, and C236,237,245S) were generated using the site-directed mutagenesis kit from NEB (New England Biolabs, Cat#: E0554S). The C236,237,245S mCherry-ZDHHC5 construct was generated using the site-directed mutagenesis kit using the C236,237S mCherry-ZDHHC5 construct as a template. Primers used to generate all constructs can be found in Supplemental Table 1.

#### Click Chemistry-Based Metabolic Labelling

We adapted the click chemistry protocol from Presolski, Hong, and Finn (2011)^33^, with modifications. HEK293T ZDHHC5^KO^ cells were seeded in 60 mm plates. At 70–80% confluence, the medium was changed to DMEM supplemented with 5% charcoal-stripped FBS. Cells were then transfected with FLAG-tagged NOD2, either the wild-type (WT) or Crohn’s disease-associated variants. At 16–20 hours post-transfection, 15-HYDA was prepared for supplementation. Briefly, in a glass vial, 2.5 µL of 120 mM KOH was added to 2.5 µL of 100 mM alkynyl palmitic acid and mixed by tapping the tube until a white solution was obtained. The mixture was then heated to 65°C for 5 minutes in a water bath, yielding a clear solution. Fatty-acid-free bovine serum albumin (FAFBSA) was dissolved in serum-free DMEM to yield a 20% (m/v) stock. Immediately after removing the lipid solution from the bath, 125 µL of 20% FAFBSA was added to the tube and mixed by pipetting until a uniform solution was obtained. The FAFBSA-lipid solution was then incubated at 37°C for 15 minutes before 125 µL of the solution was added directly to the starvation media of each 60 mm plate. Cells were treated for 4 hours before harvesting by scraping into PBS and were pelleted by centrifugation at 500 rcf for 5 minutes.

Cells were lysed in Click Chemistry Lysis Buffer (LB; 50 mM HEPES pH 7.4, 150 mM NaCl, 1% NP-40, 0.5% sodium deoxycholate, 2 mM MgCl_2_, 1 mM PMSF, and 1x EDTA-free protease inhibitor) for 15 minutes on ice, followed by centrifugation at 16,000 rcf for 10 minutes at 4°C. The supernatant was transferred to a new tube, and SDS was added to a final concentration of 0.1%. Protein concentrations were determined using the Pierce BCA assay (Thermo Scientific, Cat#: 23225, Lot#: ZL400235). FLAG affinity beads were washed twice with 1 mL of lysis buffer, after which equal amounts of protein were added to each tube of beads. Samples were nutated for 16–20 hours at 4°C to capture the FLAG-tagged NOD2.

Following capture, stock solutions of THPTA (100 mM; Vector Laboratories, Cat#: CCT-1010, Lot#: 5987A), CuSO_4_ (50 mM; Sigma-Aldrich, Cat#: 1003398912, Lot#: MKCQ4337), sodium ascorbate (300 mM; Jena Bioscience, Cat#: CLK-I005, Lot#: KLI019-004), and Cy5.5 picolyl azide (2 mM; Vector Laboratories, Cat#: CCT-1182, Lot#: 1676X) were prepared. The beads were washed three times with 500 µL of lysis buffer, and the final wash was completely removed. Reagents were then added to the beads in the following order, with thorough vortexing between each addition: 83 µL lysis buffer, 5 µL THPTA, 2 µL CuSO4, 5 µL sodium ascorbate, and 5 µL Cy5.5 picolyl azide. Tubes were then briefly centrifuged and vortexed to resuspend the beads in the reaction mixture. Tubes were incubated at 37°C for 2 hours in darkness and vortexed gently after 1 hour. Following incubation, the beads were washed three times with 500 µL of lysis buffer, and the final wash was removed completely. Protein was eluted using 1x Laemmli Sample Buffer with 100 mM DTT and resolved by SDS-PAGE. Following electrophoresis, the gel was imaged using an Azure 600 Imaging System at 700 nm to visualize the Cy5.5 signal. Proteins were then transferred from the gel to a PVDF membrane using a Trans-Blot Turbo system (Bio-Rad) and probed with an anti-FLAG antibody to visualize the amount of FLAG-NOD2 pulled down.

#### Confocal Microscopy

HCT116 cells were seeded in 12-well plates containing 18-mm circular glass coverslips and allowed to grow to 80% confluence. Cells were then transfected using 0.5 µg of GFP-NOD2 or GFP-NOD1 and 0.5 µg of mCherry-zDHHC5 or indicated truncation or mutant using X-tremeGENE HP (Lot#: 89878600) and the manufacturer’s protocol. At 16–20 hours after transfection, cells were washed with PBS (Wisent, Cat#: 311-010-CL, Lot#: 311010285) and fixed with 4% paraformaldehyde for 20 minutes at room temperature. Coverslips were wrapped in foil and protected from light during all subsequent incubation steps. Coverslips were washed with PBS and incubated in quenching-permeabilization solution containing 100 mM glycine and 0.1% (v/v) Triton X-100 for 20 minutes at room temperature. Coverslips were then blocked with 5% BSA for 1 hour at room temperature, followed by overnight incubation at 4°C with rabbit anti-FLAG (ProteinTech, Cat#: 20543-1-AP, Lot#: 00132231) diluted 1:100 in 5% BSA. Following primary antibody incubation, coverslips were washed twice with PBS before being incubated with CoraLite488-conjugated goat anti-rabbit IgG (H+L) (ProteinTech, Cat#: SA00013-2, Lot#: U0000834) diluted 1:1000 for 1 hour at room temperature. Coverslips were then washed twice with PBS and stored in PBS at 4°C until imaging.

Confocal images were acquired using an Intelligent Imaging Innovations (3i) Marianas™ spinning-disk confocal microscope built on a Zeiss Axio Observer 7 platform with Definite Focus 3. Samples were illuminated using the integrated laser lines of the LaserStack v4 (3i), specifically 488 nm (200 mW) and 561 nm (140 mW), which are delivered through single-mode optical fibers. Images were collected using a Plan-Apochromat 63×/1.4 NA oil objective (Zeiss) and captured on a Hamamatsu ORCA Fusion-BT sCMOS camera (2304×2304 pixels; 6.5 µm pixel size) controlled by SlideBook (3i). Image processing and cell membrane fluorescence intensity were quantified using the region of interest tool in ImageJ2 open-source image processing software^34^. Colocalization analysis was conducted using the BIOP JACoP plugin^35^ available on GitHub.

#### Co-Immunoprecipitations

HEK293T cells were plated in 6-well plates and transfected at 80% confluence using X-tremeGENE HP (Lot#: 89878600). Cells were transfected with 1 µg FLAG-NOD2, 1 µg soluble mCherry, 1 µg of the indicated mCherry-zDHHC5 construct, or a combination, per well. After 16–20 hours, cells were harvested in phosphate-buffered saline (PBS; Lot#: 311010285) and lysed in NP-40 buffer (50 mM Tris-HCl pH 7.5, 150 mM NaCl, 1% NP-40) containing protease inhibitors (Roche, Cat#: 11697498001, Lot#: 86108600). Lysates were incubated on ice for 1 hour with gentle vortexing at 30 minutes, then clarified by centrifugation at 14,000 rcf for 10 minutes at 4°C. Protein concentrations were determined using the Pierce BCA assay. Equal amounts of protein were incubated with RFP-Catcher agarose beads (Antibodies-Online, Cat#: ABIN5311510, Lot#: 052310) for 1 hour at 4°C with nutation to immunoprecipitate mCherry-ZDHHC5. Beads were washed three times with NP-40 lysis buffer, followed by two washes with Tris-buffered saline (TBS; 20 mM Tris-HCl pH 7.6, 150 mM NaCl), and eluted in 30 µL of 2X Laemmli sample buffer. A 2.5% total lysate input was reserved for each condition. Input and eluate samples were denatured at 37°C for 20 minutes to minimize aggregation of ZDHHC5 prior to electrophoresis. Proteins were resolved by SDS-PAGE on 10% Tris-Glycine gels and transferred to polyvinylidene fluoride (PVDF) membranes using a Bio-Rad Trans-Blot Turbo system. Membranes were blocked in 8% non-fat skim milk (NFSM) for 1 hour and incubated with mouse anti-FLAG (DSHB, Cat#: 12C6c, RRID: AB_2890618) and rabbit anti-mCherry (Cell Signaling Technology, Cat#: 43590, Lot#: 00176460) antibodies (1:1000) for 1.5 hours at room temperature in 5% NFSM. Membranes were washed five times with TBST (TBS containing 0.1% Tween-20) for 5 minutes each and incubated with fluorescent secondary antibodies (AzureSpectra goat anti-mouse 700, Cat#: AC2129, Lot#: 231214-28; goat anti-rabbit 800, Cat#: AC2134, Lot#: 240530-37) at a 1:10,000 dilution for 1 hour in 5% NFSM. Membranes were then washed five times with TBST for 5 minutes each, then visualized using the Azure 600 Imager System with 700 nm and 800 nm excitation channels.

#### Statistical Analysis

Post-acquisition micrographs were adjusted for brightness and contrast and exported as TIFF files using ImageJ2/FIJI. Plasma membrane fluorescence intensity and colocalization between FLAG-NOD2 and mCherry-zDHHC5 were quantified using region-of-interest analysis in ImageJ. Click chemistry blot band intensities were also quantified using ImageJ2/FIJI. Statistical analyses were performed using GraphPad Prism (version 11.0.2; GraphPad Software, Inc.). For microscopy-based quantifications, we determined statistical significance using one-tailed Student’s t-tests. For click chemistry blot quantifications, we determined statistical significance using a one-way repeated-measures ANOVA with Dunnett’s multiple comparisons test, comparing each condition to the NOD2-alone control lane. A significance threshold of P < 0.05 was applied throughout, and significant differences are indicated in the text and figures.

## Acknowledgments

Gregory D. Fairn holds a Tier 1 Canada Research Chair in Multiomics of Lipids and Innate Immunity. Noah R. Martin is a recipient of a graduate scholarship from the I3V Wave and the Dalhousie Medical Research Development Office. Ronan Jensen is a recipient of a graduate scholarship from the Dalhousie Medical Research Development Office. This research is supported by a Project grant from the Canadian Institutes of Health Research, PJT-198174, awarded to GDF.

## References

1. Girardin, S.E. et al. Nod2 is a general sensor of peptidoglycan through muramyl dipeptide (MDP) detection. J Biol Chem 278, 8869–8872 (2003).

2. Stafford, C.A. et al. Phosphorylation of muramyl peptides by NAGK is required for NOD2 activation. Nature 609, 590–596 (2022).

3. Caruso, R., Warner, N., Inohara, N. & Nunez, G. NOD1 and NOD2: signaling, host defense, and inflammatory disease. Immunity 41, 898–908 (2014).

4. Travassos, L.H. et al. Nod1 and Nod2 direct autophagy by recruiting ATG16L1 to the plasma membrane at the site of bacterial entry. Nat Immunol 11, 55–62 (2010).

5. Keestra-Gounder, A.M. et al. NOD1 and NOD2 signalling links ER stress with inflammation. Nature 532, 394–397 (2016).

6. Sabbah, A. et al. Activation of innate immune antiviral responses by Nod2. Nat Immunol 10, 1073–1080 (2009).

7. Hugot, J.P. et al. Association of NOD2 leucine-rich repeat variants with susceptibility to Crohn’s disease. Nature 411, 599–603 (2001).

8. Ogura, Y. et al. A frameshift mutation in NOD2 associated with susceptibility to Crohn’s disease. Nature 411, 603–606 (2001).

9. Hampe, J. et al. Association between insertion mutation in NOD2 gene and Crohn’s disease in German and British populations. Lancet 357, 1925–1928 (2001).

10. Barnich, N., Aguirre, J.E., Reinecker, H.C., Xavier, R. & Podolsky, D.K. Membrane recruitment of NOD2 in intestinal epithelial cells is essential for nuclear factor-kappaB activation in muramyl dipeptide recognition. J Cell Biol 170, 21–26 (2005).

11. Linder, M.E. & Deschenes, R.J. Palmitoylation: policing protein stability and traffic. Nat Rev Mol Cell Biol 8, 74–84 (2007).

12. Chamberlain, L.H. & Shipston, M.J. The physiology of protein S-acylation. Physiol Rev 95, 341–376 (2015).

13. Blaskovic, S., Adibekian, A., Blanc, M. & van der Goot, G.F. Mechanistic effects of protein palmitoylation and the cellular consequences thereof. Chem Phys Lipids 180, 44– 52 (2014).

14. F, S.M., et al. Mechanisms and functions of protein S-acylation. Nat Rev Mol Cell Biol 25, 488–509 (2024).

15. Rocks, O. et al. An acylation cycle regulates localization and activity of palmitoylated Ras isoforms. Science 307, 1746–1752 (2005).

16. Won, S.J., Cheung See Kit, M. & Martin, B.R. Protein depalmitoylases. Crit Rev Biochem Mol Biol 53, 83–98 (2018).

17. Lu, Y. et al. Palmitoylation of NOD1 and NOD2 is required for bacterial sensing. Science 366, 460–467 (2019).

18. Dixon, C.L. & Fairn, G.D. S-palmitoylation of NOD2 controls its localization to the plasma membrane. J Lipid Res 62, 100097 (2021).

19. Dixon, C.L., Martin, N.R., Niphakis, M.J., Cravatt, B.F. & Fairn, G.D. Attenuating ABHD17 Isoforms Augments the S-acylation and Function of NOD2 and a Subset of Crohn’s Disease-associated NOD2 Variants. Cell Mol Gastroenterol Hepatol 19, 101491 (2025).

20. Howie, J. et al. Substrate recognition by the cell surface palmitoyl transferase DHHC5. Proc Natl Acad Sci U S A 111, 17534–17539 (2014).

21. Thomas, G.M., Hayashi, T., Chiu, S.L., Chen, C.M. & Huganir, R.L. Palmitoylation by DHHC5/8 targets GRIP1 to dendritic endosomes to regulate AMPA-R trafficking. Neuron 73, 482–496 (2012).

22. Woodley, K.T. & Collins, M.O. S-acylated Golga7b stabilises DHHC5 at the plasma membrane to regulate cell adhesion. EMBO Rep 20, e47472 (2019).

23. Brigidi, G.S., Santyr, B., Shimell, J., Jovellar, B. & Bamji, S.X. Activity-regulated trafficking of the palmitoyl-acyl transferase DHHC5. Nat Commun 6, 8200 (2015).

24. Plain, F. et al. Control of protein palmitoylation by regulating substrate recruitment to a zDHHC-protein acyltransferase. Commun Biol 3, 411 (2020).

25. Mi, S. et al. AKT-mediated phosphorylation of ZDHHC5 promotes NOD1 palmitoylation and innate immune signaling. Front Immunol 17, 1819627 (2026).

26. Mi, S. et al. Metabolic orchestration of NOD1 signaling by AMPK-mediated phosphorylation of ZDHHC5. iScience 29, 115245 (2026).

27. Stauffer, T.P., Ahn, S. & Meyer, T. Receptor-induced transient reduction in plasma membrane PtdIns(4,5)P2 concentration monitored in living cells. Curr Biol 8, 343–346 (1998).

28. Dunn, K.W., Kamocka, M.M. & McDonald, J.H. A practical guide to evaluating colocalization in biological microscopy. Am J Physiol Cell Physiol 300, C723–742 (2011).

29. Ko, P.J. et al. A ZDHHC5-GOLGA7 Protein Acyltransferase Complex Promotes Nonapoptotic Cell Death. Cell Chem Biol 26, 1716–1724 e1719 (2019).

30. Rana, M.S. et al. Fatty acyl recognition and transfer by an integral membrane S-acyltransferase. Science 359 (2018).

31. Kahlson, M.A. et al. Functional dissection of the zDHHC palmitoyltransferase 5-golgin A7 palmitoylation complex. J Biol Chem 301, 110694 (2025).

32. Gok, C., Robertson, A.D. & Fuller, W. Insulin-induced palmitoylation regulates the Cardiac Na(+)/Ca(2+) exchanger NCX1. Cell Calcium 104, 102567 (2022).

33. Presolski, S.I., Hong, V.P. & Finn, M.G. Copper-Catalyzed Azide-Alkyne Click Chemistry for Bioconjugation. Curr Protoc Chem Biol 3, 153–162 (2011).

34. Rueden, C.T. et al. ImageJ2: ImageJ for the next generation of scientific image data. BMC Bioinformatics 18, 529 (2017).

35. Bolte, S. & Cordelieres, F.P. A guided tour into subcellular colocalization analysis in light microscopy. J Microsc 224, 213–232 (2006).

