## Supplemental Table 1 for "A minimal functional ZDHHC5 defines distinct requirements for NOD2 binding and *S-*acylation"

| **ZDHHC5 Large Scale Truncations** | **Primer Sequence** |
| --- | --- |
| Forward (Same forward primer for all) | GTCAAGCTTTCATGCCCGCAGAGTCTGGA |
| Reverse 527X | GACGGGCCCCTCGGTGTGTTGGGCCAGT |
| Reverse 335X | GACGGGCCCAGTAGCCAGGCCGAGGTG |
| Reverse 275X | GACGGGCCCCTGAAACTTCTGGTCGAAGGAAGGG |
| Reverse 219X | GACGGGCCCATTGGTTGTGCGTCCCCT |

| **ZDHHC5 Small Scale Truncations** | **Primer Sequence** |
| --- | --- |
| Forward (Same forward primer for all) | CTCAGATCTCGAATGCCC |
| Reverse 261X | GTAAAGCTTTTATTACTTCTCTTTCTTTGGTCTCCC |
| Reverse 251X | GTAAAGCTTTTATTAGGGTGCTGGAGAACT |
| Reverse 241X | GTAAAGCTTTTATTAGACATTGTTACAGCAGCC |
| Reverse 231X | GTAAAGCTTTTATTAGTTCACACCTCCCCG |

| **ZDHHC5 Site Directed Mutagenesis** | **Primer Sequence** |
| --- | --- |
| C245S Forward | CCGTGTTCTCAGCAGTTCTCCAG |
| C245S Reverse | CTGACATTGTTACAGCAGC |
| C236,237S Forward | CACCAATGGCAGCAGTAACAATGTCA |
| C236,237S Reverse | AAGGGGTTCACACCTC |
| C134S Forward | TGATCATCACAGCCCCTGGGTGA |
| C134S Reverse | AATTCCTCCACACAGTTGTCAC |
